# Injectable microcapillary network hydrogels engineered by liquid-liquid phase separation for stem cell transplantation

**DOI:** 10.1101/2023.07.22.550127

**Authors:** Akihiro Nishiguchi, Shima Ito, Kazuhiro Nagasaka, Hiyori Komatsu, Koichiro Uto, Tetsushi Taguchi

## Abstract

Injectable hydrogels are promising carriers for cell delivery in regenerative medicine. However, injectable hydrogels composed of crosslinked polymer networks are often non porous and prevent biological communication with host tissues through signals, nutrients, oxygen, and cells, thereby limiting graft survival and tissue integration. Here we report injectable hydrogels with liquid-liquid phase separation-induced microcapillary networks (µCN) as stem cell-delivering scaffolds. The molecular modification of gelatin with hydrogen bonding moieties induced liquid-liquid phase separation when mixed with unmodified gelatin to form µCN structures in the hydrogels. Through spatiotemporally controlled covalent crosslinking and dissolution processes, porous µCN structures were formed in the hydrogels, which can enhance mass transport and cellular activity. The encapsulation of cells with injectable µCN hydrogels improved cellular adhesion, spreading, migration, and proliferation. Transplantation of mesenchymal stem cells with injectable µCN hydrogels enhanced graft survival and recovered hindlimb ischemia by enhancing material-tissue communication with biological signals and cells through µCN. This facile approach may serve as an advanced scaffold for improving stem cell transplantation therapies in regenerative medicine.

## Introduction

Stem cell transplantation therapy holds immense potential to treat intractable diseases including ischemia,^1^ myocardial infarction,^2^ and spinal cord injury.^3^ Despite the substantial advances in regenerative medicine, current approaches rely on direct injection of cell suspension, which often results in poor delivery efficiency, cell survival, migration, proliferation, and integration with host tissue.^4, 5^ Infused cells diffuse and accumulate in organs where their accumulation is undesirable (e.g., in the lungs and liver when intravenously infused^6^) and undergo apoptotic cell death.^7^ Injectable hydrogels that form gels in the body have been used as cell-delivering biomaterials, demonstrating local delivery and retention of cells by adhering to target tissues.^8^ Additionally, injectable hydrogels can function not only as reservoirs of cells but also as scaffolds to enhance the regenerative capacity of stem cells.^9^ Although substantial efforts have been focused on designing cell-delivering injectable hydrogels, including implementing covalent crosslinks,^10^ cell-polymer hybrids,^11^ inclusion complexes,^12^ and protein hydrogels,^13^ their major limitation is lack of macro/microporous structures to exchange biological signals and cells with host tissues. Typically, injectable hydrogels entrap transplanted cells in densely crosslinked polymer networks (e.g., the pore size of a poly(ethylene glycol) hydrogel is approximately 20 nm^14^), which prevents the diffusion of nutrients and oxygen. These crosslinked hydrogels (non-porous hydrogels) may restrict the survival of transplanted cells and the cellular infiltration necessary for rapid integration.^15^ Thus, injectable hydrogels with macro/micropores have been studied to improve graft survival. However, the current approaches for fabricating porous injectable hydrogels using particle porogens,^16–18^ microgels,^19, 20^ and peptide nanofibers,^21^ fail to provide interconnected pores and void spaces to induce cell infiltration. Clinically available fibrin gels have been used for cell delivery,^22^ but the porosity was estimated to be less than 1 µm at a clinically used concentration (40 mg/mL), which was too small for cell infiltration.^23^ These drawbacks may lead to poor host integration, especially for vascularization.^24, 25^

Liquid-liquid phase separation (LLPS) is often seen in biological systems such as intracellular membraneless organelles and mussel byssal plaques, where it plays a central role in organelle formation, biomolecular condensation, and signal transduction.^26^ In response to damage and aging, reversible assembly of proteins forms LLPS for DNA damage repair,^27^ while irreversible morphological changes cause solid aggregates in the LLPS, leading to diseases.^28^ These dynamic LLPS processes are regulated by post translational modifications and multivalent molecular interactions in intrinsically disordered protein regions (IDRs).^26^ IDRs can function as a self-associating, sticky patch through non-covalent interactions such as hydrogen bonding, and electrostatic or hydrophobic interactions, which assemble proteins to induce LLPS. For example, hyperphosphorylation in IDRs of a nucleolar phosphoprotein, Ki-67, drove LLPS through an imbalance of electrostatic interactions.^29^ Moreover, in mussel foot proteins, a post translationally modified amino acid, *L*−3,4-dihydroxyphenylalanine (Dopa), mediates hydrogen bonding interaction and undergoes LLPS for wet adhesion.^30^ Inspired by these biological systems, we assumed that LLPS behaviors can be regulated by molecular modification with functional groups and their engineered structures can be used as a porogen to design cell-delivering injectable porous hydrogels. Here we report the use of injectable hydrogels with LLPS-generated microcapillary networks (µCN) to regulate cellular activity and improve stem cell transplantation therapy through enhanced material-tissue communications. The extracellular matrix-derived protein gelatin was chemically modified with ureidopyrimidinone (UPy) units.^31, 32^ Gelatin modified with UPy units (GUPy) formed LLPS structures by mixing with non-modified gelatin via hydrogen bonding-mediated self-association. Injectable µCN hydrogels were engineered through mild Michael addition-based crosslinking of the gelatin matrix and dissolution of GUPy. µCN hydrogels promote cellular adhesion, spreading, migration, and proliferation by improving mass transport and providing adhesive scaffolds. µCN hydrogels improved graft survival of mesenchymal stem cells (MSCs) and therapeutic efficacy for hindlimb ischemia. This facile approach can be used to design injectable hydrogels with controlled internal structures that may improve the therapeutic efficacy of stem cell transplantation.

## Results

### LLPS-induced µCN structures using self-associating gelatin

We previously reported GUPy-based supramolecularly crosslinked hydrogels as tissue adhesive.^33^ Here, we found that mixing skin-derived gelatin (sG) solution with tendon derived gelatin modified with UPy units (tGUPy) solution formed LLPS structures, although both were gelatin-based polymers (Fig. 1a). Because hydrogen bonding is a key regulator of the LLPS of proteins,^30^ molecular modification of gelatin with UPy units that function as self-associating patches via non-covalent quadruple hydrogen bonding is expected to cause LLPS in sG and tGUPy. For the visualization of the LLPS structures, sG and tGUPy were dissolved in phosphate-buffered saline (PBS), respectively, and mixed by applying shear and cooled to room temperature (below the sol-gel transition temperature) to form hydrogels. Interestingly, micrometer-scale fiber-like structures were observed in the hydrogels when sG and tGUPy were mixed at equal volumes (Fig. 1b). Confocal laser scanning microscopy (CLSM) observations revealed that the mixture of sG+tGUPy formed bicontinuous phase-separated structures comprising fiber-like tGUPy and sG as a matrix and increased the turbidity of the hydrogels, indicating the formation of LLPS (Fig. 1c,d). Fiber-like LLPS structures were distributed homogeneously throughout the hydrogels, and sG and tGUPy were separately localized in the hydrogels, indicating the formation of segregated LLPS (Fig. 1e,f). The fiber-like structures of tGUPy were connected to each other and distributed three dimensionally in the hydrogels. To explore the mechanism of LLPS formation, phase diagrams of sG and tGUPy were constructed (Fig. 1g). Typically, phase-separated structures in LLPS have been characterized as droplets and bicontinuous structures.^34^ In this unique segregative LLPS system, we observed the droplets and two bicontinuous structures of tGUPy: gyroid-like structures seen in the non-solvent phase separation of *block*-copolymers^35^ and thinner fibrous structures with capillary networks a few micrometers in diameter, which we defined as gyroid and µCN, respectively. µCN structures were observed at high concentrations of tGUPy and sG, and showed longer tube lengths and thinner diameters (Fig. 1h). At low sG concentrations, thick gyroid structures with short tube lengths were formed. Droplets were observed in the sG matrix at low tGUPy concentrations. Among the LLPS structures, µCN structures displayed a blood capillary network-like morphology, which is useful as a porogen for cell encapsulation.

**Fig. 1.**
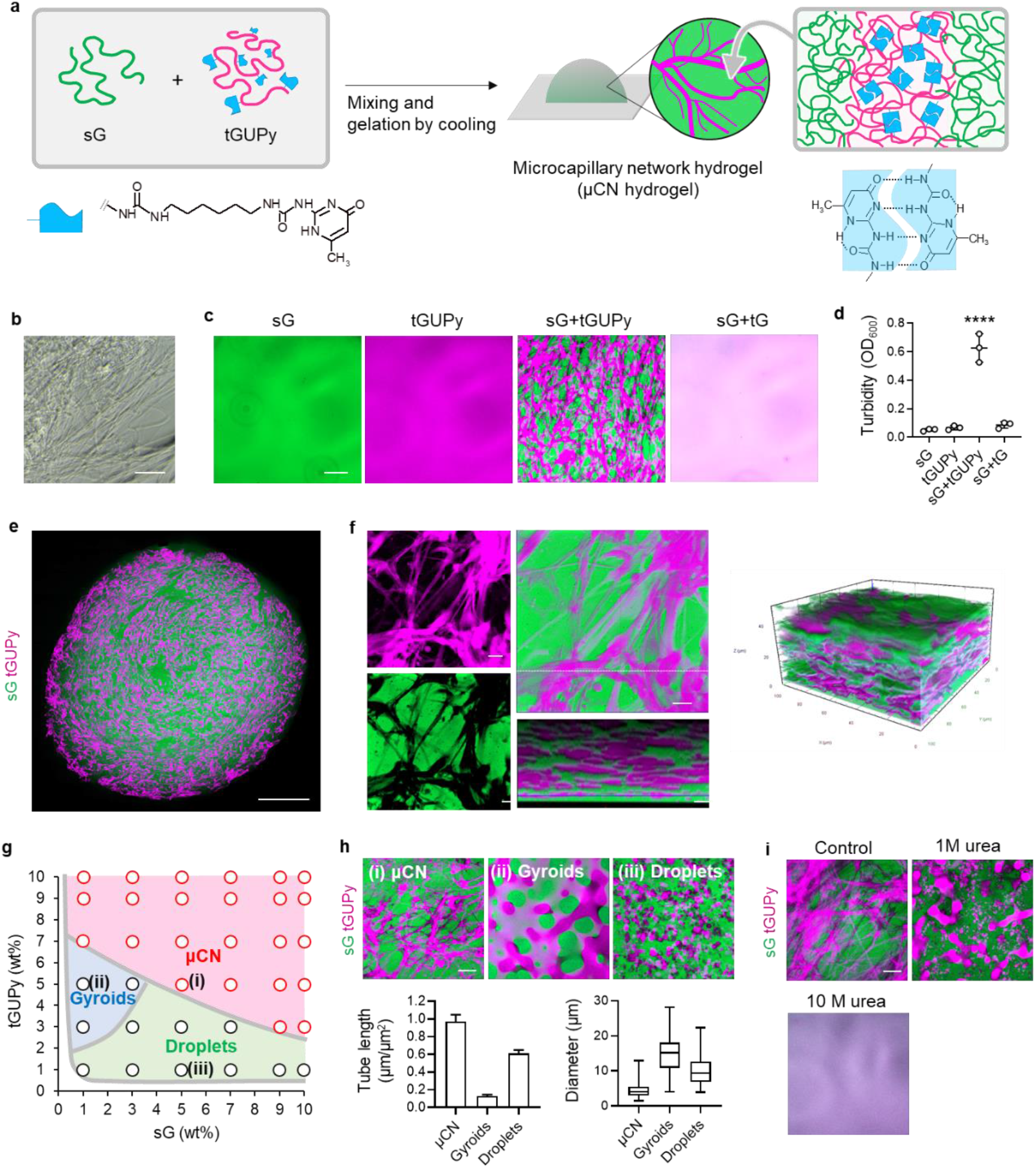
LLPS-induced µCN structures in hydrogels. **a**, Schematic illustration of the preparation of microcapillary networks (µCN) hydrogels through liquid-liquid phase separation (LLPS). Skin-derived gelatin (sG) and tendon-derived gelatin modified with the ureidopyrimidinone (UPy) unit (tGUPy) were mixed and shear force was applied, resulting in the formation of hydrogels with LLPS structures after cooling to room temperature. **b**, Phase contrast image of µCN hydrogels. Fibrous capillary network structures were observed. **c,d**, CLSM images and turbidity (OD600) of hydrogels composed of sG, tGUPy, sG+tGUPy, and sG+tG. sG and tGUPy were fluorescently labeled with fluorescein (green) and Cy5.5 (violet). **e,f**, Whole CLSM and reconstructed CLSM images of µCN hydrogels of sG+tGUPy. **g**, Phase diagram of LLPS structures of sG and tGUPy. **h**, CLSM images of hydrogels of µCN (sG+tGUPy = 1:1), gyroids (sG+tGUPy = 1:5), and droplets (sG+tGUPy = 5:1) and their quantification of tube length and diameter. The images of (i)-(iii) correspond with the preparation conditions shown in (g). **i**, The blocking effects of hydrogen bonding on LLPS structures with 1 and 10 M urea. \*\*\*\**P* < 0.0001, analyzed by one-way analysis of variance (ANOVA) with Tukey’s multiple comparison *post hoc* test. Scale bars represent 10 µm for (f), 50 µm for (b,c,h,i), and 1 mm for (e).

To understand the mechanism of formation of µCN structures, the effects of the molecular weight of sG, degree of substitution (D.S.) of UPy units, and derivation of gelatin (skin or tendon) on LLPS formation were evaluated. sG of more than 60 kDa and 42% D.S. of UPy units were required for the formation of µCN structures (Supplementary Fig. 1a,b). These results indicate that the high molecular weight of sG and high D.S. of the UPy units accelerated segregative LLPS due to increased incompatibility. Regarding the derivation of gelatin, the combination of sG and tGUPy formed µCN structures, while other combinations formed dense networks (tG+tGUPy) or thick gyroid structures (sG+sGUPy, tG+sGUPy) (Supplementary Fig. 1c). Owing to the higher molecular weight of tG than sG and the difference in amino acid sequences, tGUPy showed strong self associating properties via intermolecular hydrogen bonding between not only UPy-UPy but also tG-tG, forming µCN-type LLPS structures in the hydrogels. Additionally, tGUPy formed µCN structures when mixed with bovine serum albumin (BSA), indicating that this approach can be extended to various proteins to engineer porous protein hydrogels (Supplementary Fig. 1d). These results suggest that self-associating gelatin is a powerful tool for designing hydrogels with controlled internal structures.

To address the effect of hydrogen bonding on LLPS, blocking experiments were performed using urea. CLSM observation showed that adding urea to hydrogels prevent LLPS formation (Fig. 1i). These results highlight the importance of multivalent hydrogen bonding between the UPy units in gelatin for inducing LLPS. Moreover, when LLPS solution was incubated at 37 °C in which gelation did not occur, LLPS solution underwent bulk macroscale phase separation containing droplet-rich phase in upper solution and capillary-rich phase in lower solution (Supplementary Fig. 2). This result indicates that the LLPS structures were transiently formed when mixed by applying shear stress and were maintained by the sequential gelation of gelatin.

### Engineering covalently crosslinked µCN hydrogels

Two processes are required to use injectable µCN hydrogels as cell scaffolds: covalent crosslinking of the matrix components (sG) and dissolution of the LLPS structures (tGUPy). Michael addition between the thiol and vinyl sulfone groups was used to crosslink the sG matrix (Fig. 2a). Mild covalent crosslinking enabled the encapsulation of cells in µCN hydrogels and provided mechanical strength for tissue adhesion. After gelation, hydrogels were immersed in PBS to dissolve tGUPy and form porous µCN structures in hydrogels. Notably, the modification of sG with thiol and vinyl sulfone groups did not affect LLPS formation with tGUPy. sG was chemically modified with thiol (sGTH) and vinyl sulfone groups (sGVS) at a defined D.S. (Supplementary Table 1). The introduction of functional groups was determined by ^1^H-NMR spectroscopy (Supplementary Fig. 3). The gelation of sGTH and sGVS was confirmed by rheological measurements, and the gelation kinetics and mechanical strength depended on the D.S. of the thiol and vinyl sulfone groups (Supplementary Fig. 4). Non-porous and µCN hydrogels were prepared by mixing sGTH, sGVS, and tG or tGUPy (10 wt%, volume ratio; 1:1:2). Rheological measurements showed that µCN hydrogels reached a gelation plateau within 20 min and showed faster gelation kinetics than the non-porous hydrogels (Supplementary Fig. 5). In addition, µCN hydrogels exhibited higher fracture strain and stress than the non-porous hydrogels, indicating improved mechanical strength (Fig. 2b). The strength of adhesion to the collagen membrane was enhanced by LLPS formation (Fig. 2c). LLPS may increase the local polymer concentration of sGTH and sGVS in the matrix and improve the mechanical strength, which is consistent with a previous report.^36^

**Fig. 2.**
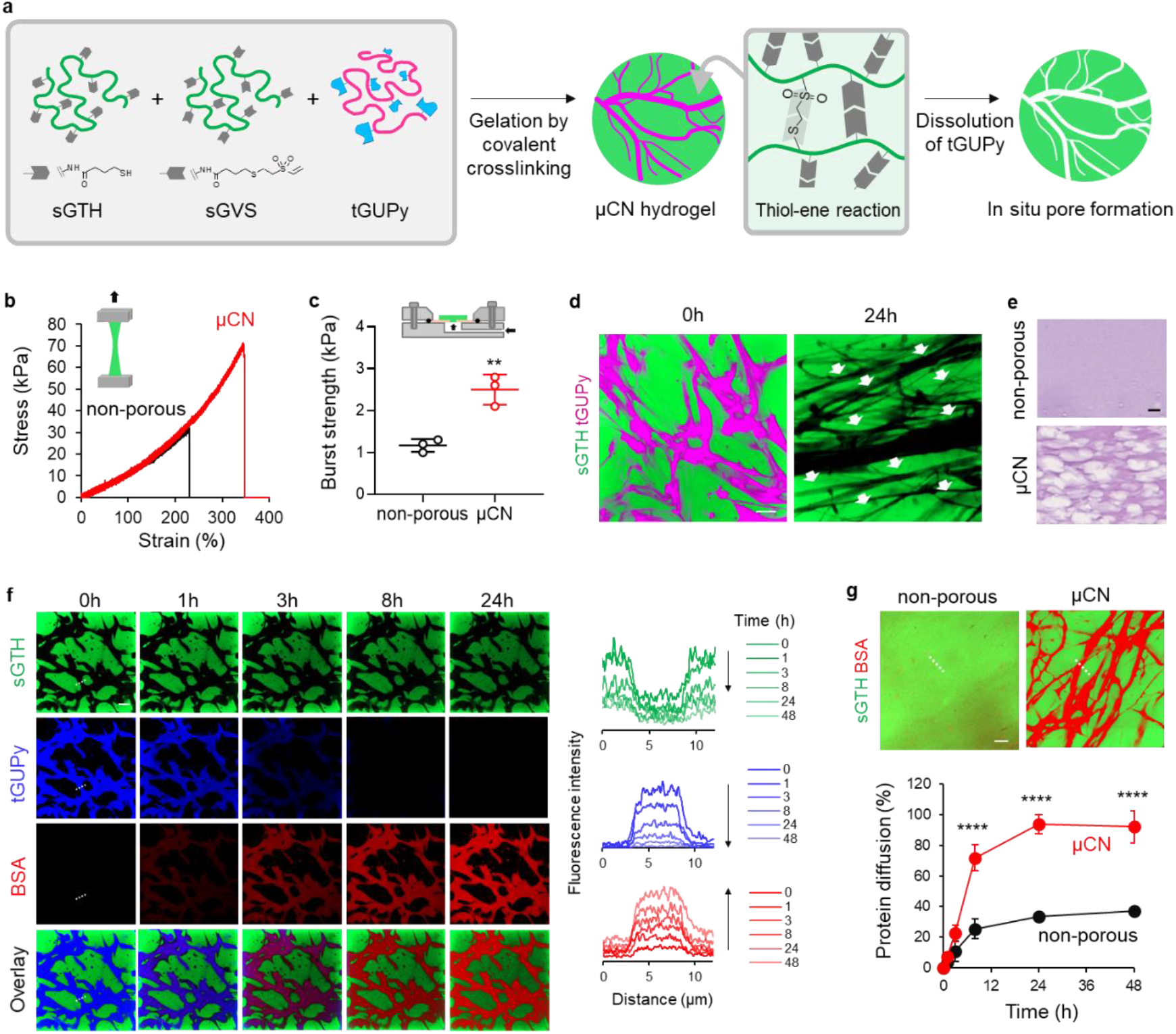
Engineering μCN hydrogels through spatiotemporally controlled covalent crosslinking and dissolution process. **a**, µCN hydrogels were formed by thiol-ene covalent crosslinking of sG matrix. tGUPy was dissolved under physiological conditions and media diffused into µCN structures. **b,c**, Tensile test and adhesion test of non-porous and µCN hydrogels (n=3). The hydrogels were used before swelling. **d**, CLSM images of µCN hydrogels before and after immersion in PBS. The arrows denote µCN structures. sGTH-fluorescein (green) and tGUPy-Cy5.5 (violet) were used. **e**, HE images of non porous and µCN hydrogels before and after the immersion in PBS. **f**, Diffusion test of proteins into µCN hydrogels. Rhodamine-labeled BSA was added to µCN hydrogels of sGTH-fluoresceine (green) and tGUPy-Cy5.5 (blue). Line-scan profiles showed the change of fluorescence intensity during the incubation. **g**, Comparison of BSA diffusion between non-porous and µCN hydrogels after 48 h of incubation (n=3). Protein diffusion was quantified from CLSM line scan profiles and the fluorescence intensity of original solution of rhodamine-labeled BSA was set to 100%. Data are presented as the mean ± s.d. from a representative experiment (biologically independent samples). \*\**P* < 0.01, \*\*\*\**P* < 0.0001 analyzed by the two-tailed Student’s *t*-test (c) and two-way ANOVA with Tukey’s multiple comparison *post hoc* test (g). Scale bars represent 10 µm.

CLSM observations revealed that tGUPy disappeared during incubation in the media and formed porous µCN structures in the hydrogels, indicating that tGUPy dissolved under physiological conditions and functioned as a porogen (Fig. 2d). Cross-sectional images stained with hematoxylin & eosin (HE) showed that non-porous hydrogels possessed dense solid-like structures without micropores, while µCN hydrogels formed highly porous structures (Fig. 2e). The swelling ratios of the non-porous and µCN hydrogels after immersion in PBS for 24 h were 32.1 and 25.4%, respectively, and their elastic moduli were similar (Supplementary Fig. 6). The weight of µCN hydrogels reduced to 65% after 24 h in PBS, and the dissolution of tGUPy was confirmed (Supplementary Fig. 7). The gelatin-based µCN hydrogels were enzymatically degraded by collagenase, indicating their biodegradability.

To determine whether this spatiotemporal approach to control the inner structures of hydrogels could enhance mass transport through µCN structures, protein diffusion into the hydrogels was monitored. CLSM observations and line-scan profiles revealed that BSA diffused into µCN structures during incubation (Fig. 2f). µCN hydrogels achieved over 70% diffusion of BSA within 8 h and showed higher mass transport than non-porous hydrogels (Fig. 2g). Moreover, µCN hydrogels showed improved transport of glucose, a key nutrient for cell survival, compared to non-porous hydrogels (Supplementary Fig. 8). µCN hydrogels enabled the diffusion of proteins through micrometer-scale capillaries, similar to vasculatures. In the body, the maximum distance between cells and blood vessels that still enables the supply of oxygen and nutrients is 150–200 µm.^37^ Hydrogels thicker than 200 µm limit diffusion, causing hypoxia and necrosis.^38^ Engineered porous µCN structures were three-dimensionally distributed at high density in the hydrogels, which may support mass transport to maintain the viability and functions of transplanted cells.

### Cellular behavior in µCN hydrogels

To address the effects of µCN structures on cellular activity, mouse myoblasts were encapsulated in µCN hydrogels and their cellular behavior was analyzed. To improve the therapeutic efficacy of cell transplantation, it is crucial to design injectable hydrogels with physicochemical cues that enhance cellular functions. The cells were mixed with sGTH, sGVS, and tGUPy solutions (10 wt%, volume ratio: 1:1:2) for encapsulation (Fig. 3a). During culture, µCN pores were formed, and the media diffused into the hydrogels. The cytocompatibility of sGTH and sGVS was confirmed using a cell viability assay using mouse fibroblasts (Supplementary Fig. 9). CLSM observation revealed that cells in non porous hydrogels possessed round and poor adherent morphology, while µCN hydrogels drastically improved cellular adhesion and spreading (Fig. 3b). To understand how cells spread in the hydrogels, the localization of cells and µCN structures was observed by CLSM using sGTH-fluorescein. Although non-porous hydrogels restricted cell spreading through crosslinked polymer networks, cells in µCN hydrogels showed highly elongated morphologies along the porous µCN structures (Fig. 3c). In non-porous hydrogels, cells partially adhere to the hydrogels through cell-adhesive ligands (e.g., RGD sequences), but cell spreading depends on slow enzymatic degradation of the sG matrix. The µCN hydrogels rapidly formed porous µCN structures and accelerated cellular adhesion and spreading.

**Fig. 3.**
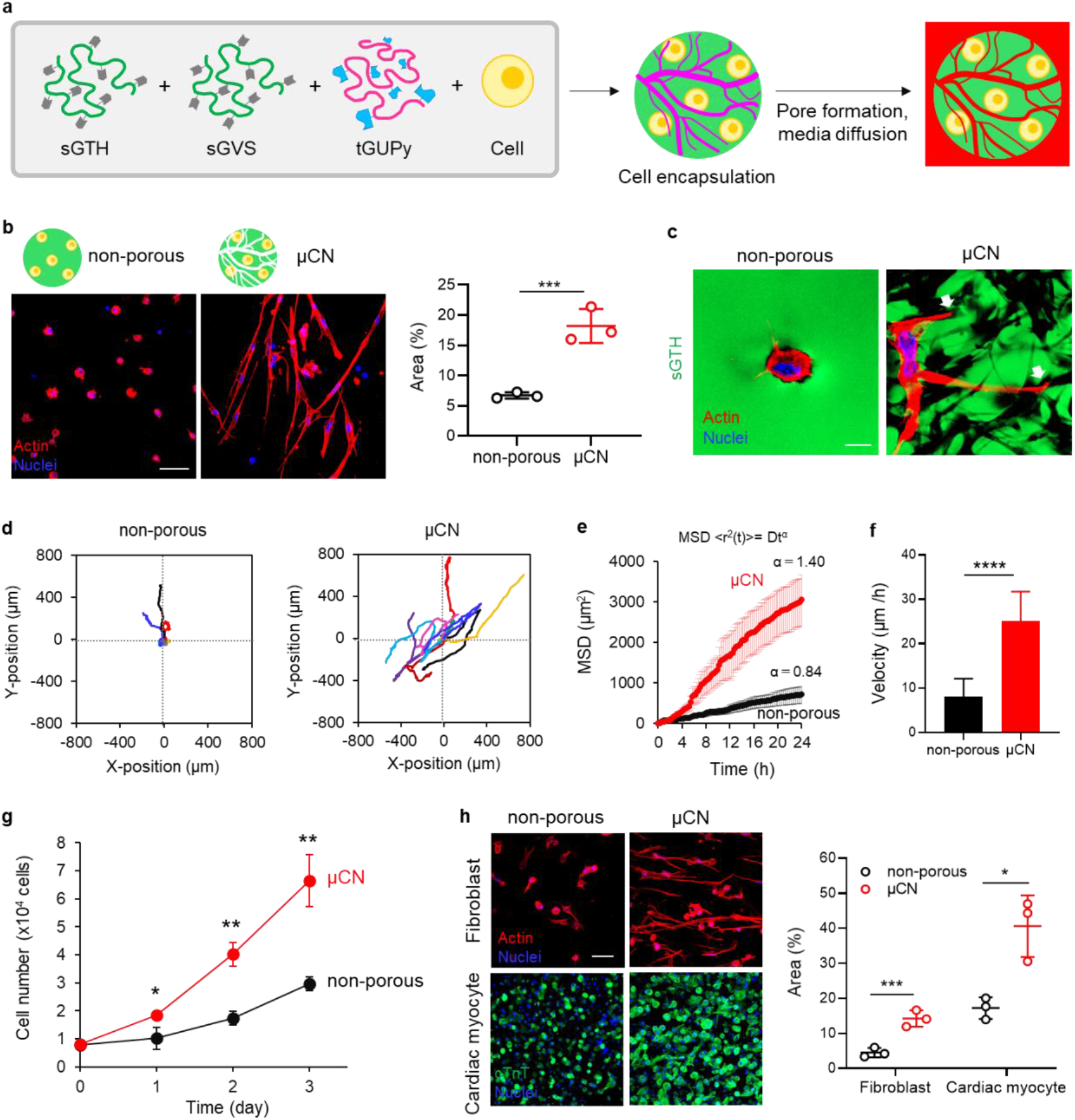
Enhanced cellular adhesion, spreading, migration, and proliferation in µCN hydrogels. **a**, Schematic illustration of cell encapsulation in μCN hydrogels. Mouse myoblast (C2C12 cells) were encapsulated and tGUPy was dissolved and exchanged with media at 37 °C. **b**, CLSM images of cells encapsulated in non-porous and µCN hydrogels for 24 h. Actin and nuclei were stained with phalloidin (red) and DAPI (blue). Cells were scanned in 50 µm thickness and the obtained images were overlaid. The adhesion area of cells in the hydrogels was quantified (n=3). **c**, Magnified CLSM images of cells encapsulated in non-porous and µCN hydrogels for 24 h. sGTH-fluorescein (green), phalloidin (red), and DAPI (blue) were used for visualization. **d**, Trajectories of GFP expressing cells cultured in hydrogels during 24 h of incubation. The trajectories were plotted at a common origin for visualization. **e**, Mean-square displacements (MSD) was calculated from trajectories shown in d. **f**, Velocity of cells in hydrogels. **g**, Proliferation tests of cells in non-porous and µCN hydrogels (n = 3). **h**, CLSM images of encapsulation of fibroblasts (normal human dermal fibroblast, NHDF) and iPS-derived cardiac myoblasts-NHDF in non-porous and µCN hydrogels. Cells were stained with phalloidin (red) or cTnT antibody (green) and DAPI (blue). Data are presented as the mean ± s.d. from a representative experiment (biologically independent samples). \**P* < 0.05 \*\**P* < 0.01, \*\*\**P* < 0.001, \*\*\*\**P* < 0.0001, analyzed by the two-tailed Student’s *t*-test. Scale bars represent 50 µm for (b,h) and 10 µm for (c).

The trajectories of GFP-expressing myoblast cells in µCN hydrogels promoted cellular migration compared to those in non-porous hydrogels (Fig. 3d, Supplementary Movie 1). The value of α, the exponent of mean-square displacement (MSD) of µCN hydrogels, exceeded 1, and cellular migration appeared super-diffusive, typically representing persistent random migration,^39^ owing to locally oriented µCN structures (Fig. 3e). The velocity of the cells in µCN hydrogels was higher than that in the non-porous hydrogels (Fig. 3f). Importantly, µCN hydrogels promoted cellular proliferation and showed a 2.3-fold increase in cell number after 3 d of incubation compared to non-porous hydrogels (Fig. 3g). These results suggest that µCN structures function not only as capillaries for mass transport, but also as voids to enhance cellular adhesion, spreading, migration, and proliferation. Additionally, fibroblasts and cardiac myocytes encapsulated in µCN hydrogels showed cellular spreading and partially synchronous beating, respectively, but not in non-porous hydrogels (Fig. 3h). These results indicate that µCN hydrogels provide suitable microenvironments for cellular adhesion and can be used in a variety of cell transplantation therapies.

### Cellular functions of MSCs in µCN hydrogels

MSCs have attracted considerable attention as a therapeutic cell source for stem cell transplantation owing to their self-renewal capacity, multilineage differentiation potential, low immunogenicity, paracrine angiogenic signaling, and immunomodulatory functions.^40^ Through the secretion of growth factors, cytokines, and extracellular vesicles, MSCs demonstrate regenerative effects against various diseases in clinical trials.^41^ We addressed the encapsulation of MSCs in hydrogels and explored how the geometry resulting from the porous structures affects cellular behavior. MSCs were encapsulated in non-porous hydrogels and three LLPS-induced porous hydrogels (µCN, gyroids, and droplets). CLSM observations revealed that µCN hydrogels substantially improved the adhesion and spreading of MSCs compared to non-porous, gyroids, and droplets (Fig. 4a). Gyroids with thick fibers surrounded MSCs, restricting their spreading, and some MSCs dropped out through voids larger than the cells. Droplets without interconnected networks entrapped MSCs in the voids and inhibited their spread. These results suggest that controlling the internal structures of hydrogels, such as pore geometry and size, using an LLPS-based approach can regulate cellular behavior. When the seeding cell number and incubation time were increased (1.5 × 10^5^ cells/20 µL, 2 d), MSCs were well distributed throughout the µCN hydrogels, exhibiting concentric local orientation morphology and distributed three-dimensionally (Fig. 4b). During the preparation of µCN hydrogels on a substrate, the pre-gel solution was subjected to shear stress, leading to the orientation of the µCN structures in the hydrogels, especially at the edge of the hydrogels, which induced MSC orientation. MSCs were oriented even at low cell density conditions (1 × 10^4^ cells/20 µL), indicating that the cell orientation originated from cell-material interaction rather than cell-cell interaction and cell crowding (Supplementary Fig. 10).

**Fig. 4.**
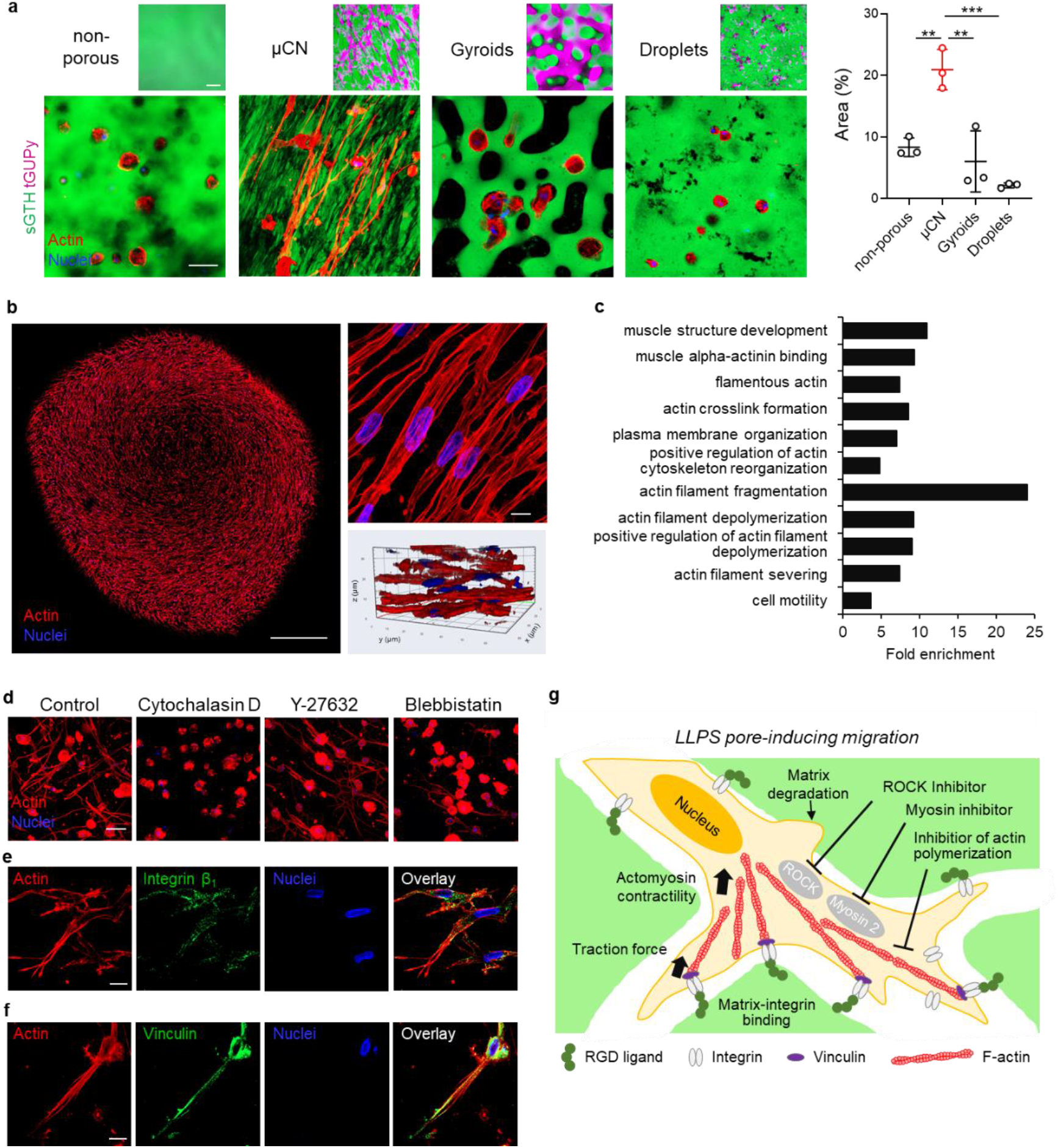
|Encapsulation of MSCs in µCN hydrogels and their angiogenic property. **a**, CLSM images of MSCs; 5 × 10^4^ cells) cultured in non-porous, μCN, gyroid, and droplet hydrogels for 24h. Hydrogels with µCN, gyroids, and droplets were prepared using sGTH+sGVS+tGUPy (1:1:2), sGTH+sGVS+sGUPy (1:1:2), and sGTH+sGVS+tGUPy (1:1:0.4). sGTH-fluorescein (green) and tGUPy-Cy5.5 (violet) were used. Actin and nuclei were stained with phalloidin (red) and DAPI (blue). **b**, Gross, magnified and three dimensional (3D-) reconstructed images of MSCs (1.5 × 10^5^ cells) cultured in µCN hydrogels for 48 h. **c**, GO analysis of MSCs cultured in non-porous and µCN hydrogels obtained from the results of RNA-seq analysis. **d**, Inhibitory tests of MSCs using cytochalasin D (10 µM), Y-27632 (10 µM), and blebbistatin (10 µM). **e,f**, CLSM images of integrin β1 and vinculin stained MSCs in µCN hydrogels. **g**, Schematic of cellular adhesion in µCN hydrogels. Data are presented as the mean ± s.d. from a representative experiment (biologically independent samples). \**P* < 0.05, \*\*\**P* < 0.001, analyzed by the two-tailed Student’s *t*-test (h) and one-way ANOVA with Tukey’s multiple comparison *post hoc* test (a). Scale bars represent 1 mm for (b [left]), 50 µm for (a,d) and 10 µm for (b [right], e, f).

To understand the effect of µCN structures on MSC function, Gene Ontology (GO) analysis was performed based on the results of the RNA-seq analysis (Fig. 4c and Supplementary Fig. 11). Several genes related to actin filaments and plasma membrane organization were specifically upregulated in µCN hydrogels. Moreover, actin filament fragmentation and depolymerization were enriched in µCN hydrogels. These results were consistent with the results of cellular adhesion, spreading, and migration seen in CLSM observations, indicating that µCN hydrogels served as scaffolds that provided a biochemical and mechanostructural microenvironment for cellular adhesion and enhanced cytoskeleton organization and cell motility. We further investigated the mechanism of cellular adhesion using small molecule inhibitors critically related to the mechanotransduction pathway. The inhibition of actin polymerization using cytochalasin D substantially suppressed MSC spreading, highlighting the important role of actin polymerization in cell spreading in µCN hydrogels (Fig. 4d). Inhibition of Rho-associated protein kinase with Y-27632 and myosin II with blebbistatin resulted in less cell spreading, indicating that cell spreading in the hydrogels was mediated by actomyosin-based contractility.^42^ Additionally, immunofluorescence staining of MSCs with an antibody against integrin β_1_ and vinculin which are a key marker of cell-extracellular matrix interaction in mechanosensing, showed that integrin β_1_ and vinculin were localized and clustered at the periphery of cells (Fig. 4e,f). Taken together, µCN hydrogels demonstrated mechanotransduction in MSCs based on cell-matrix interactions between integrin and adhesive ligands (RGD sequences), and induced spreading and migration via LLPS pores rather than matrix degradation-dependent migration (Fig. 4g).

### Treatment of hindlimb ischemia using MSC-encapsulated µCN hydrogels

MSCs are a potent medical intervention for ischemic diseases owing to their ability to secrete angiogenic factors such as vascular endothelial growth factor (VEGF). MSCs encapsulated in non-porous and µCN hydrogels produced large amounts of VEGFs at almost the same level during the 7 d of incubation (Fig. 5a). In contrast, co-culture experiments with human umbilical vein endothelial cells (HUVECs) and MSCs in hydrogels showed that µCN hydrogels enhanced the cellular adhesion and spreading of HUVECs (Fig. 5b). These results indicate that MSC-encapsulating µCN structures with similar diameter to blood capillaries can support the organization of blood vessels. Since it has been reported that the geometric control of vascular structures (aligned channels) can enhance tissue integration,^43^ µCN structures may contribute to promoting angiogenesis in host tissues. Besides VEGF secretion, it was confirmed that MSCs encapsulated in hydrogels suppressed the secretion of inflammatory cytokines, tumor necrosis factor-α (TNF-α), from macrophages through paracrine signaling compared to control and 2D-cultured MSCs (Supplementary Fig. 12). This enhanced anti inflammatory function of the hydrogels may facilitate tissue regeneration. Taken together, the combination of paracrine signals by MSCs and 3D-geometric effect by µCN hydrogels would be effective in inducing angiogenesis in host tissues and treating ischemia.

**Fig. 5.**
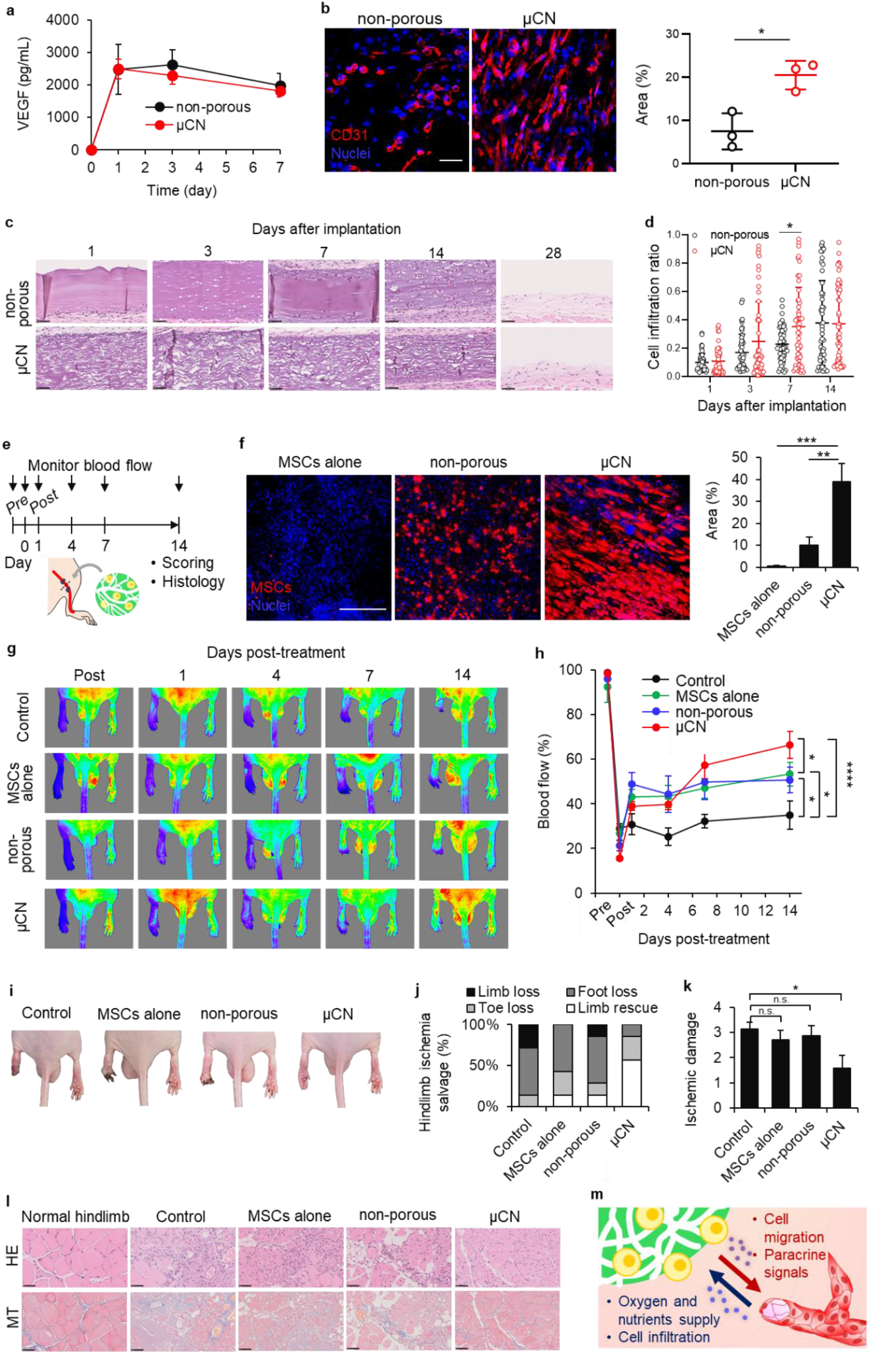
MSC-encapsulated µCN hydrogels recovered hindlimb ischemia. **a**, Secretion of VEGF from MSCs cultured in non-porous and μCN hydrogels (n=3). **b**, CLSM images and adhesion area of HUVECs and HUVECs+MSCs cultured in non-porous and µCN hydrogels (n=3). **c**, Histological observation of HE-stained images of non-porous and µCN hydrogels. **d**, Cell infiltration ratio quantified from HE images. **e**, Schematic illustration of the procedure for hindlimb ischemia mouse model. Femoral artery in left hindlimb of BALB/c nude mice was ligated and cut to induce ischemia followed by the treatment with MSCs alone or MSC-encapsulated non-porous and µCN hydrogels. **f**, Graft survival of transplanated MSCs (n=3). MSCs were stained with DiI before the injection. At day 3 after the transplantation, tissues were collected and observed by CLSM. **g**, Hindlimb blood flow monitored using a laser Doppler imaging system. **h**, Quantification of hindlimb blood flow (n=7). Blood flow denotes the percentage compared to non-treated hindlimb (right). **i**, Representative photographs of hindlimb at day 14 after the treatment. **j**, Physiological status of ischemic hindlimb at day 14. **k**, Ischemic damage of limbs were semiquantified by ischemia scores. **l**, Histological observation of HE and MT-stained cross-sectioned images of hindlimb. **m**, Schematic of the mechanism to recover ischemia by µCN hydrogels. Data are presented as the mean ± s.d. (a,b) or ± s.e.m. (d,h,k) from a representative experiment (biologically independent samples). \**P* < 0.05, \*\**P* < 0.01, \*\*\**P* < 0.001, \*\*\*\**P* < 0.0001, analyzed by the two tailed Student *t*-test (b,d) or one-way (k) and two-way (h) ANOVA with Tukey’s multiple comparison post hoc test. n.s. denotes not significant. Scale bars represent 50 µm.

Next, to evaluate cell infiltration into the hydrogels and their biodegradability and biocompatibility, the hydrogels were subcutaneously implanted into mice. Histological observation showed that porous µCN structures were formed in µCN hydrogels on day 1 after transplantation, and cells initiated infiltration (Fig. 5c). µCN hydrogels promoted cell infiltration from host tissues through µCN structures (Fig. 5d). On day 28, the subcutaneously implanted hydrogels had degraded.

Finally, we evaluated the therapeutic efficacy of MSC-encapsulated hydrogels in a mouse model of hindlimb ischemia. Critical hindlimb ischemia is a peripheral artery disease that causes necrosis and limb loss. There are numerous studies on the treatment of ischemia using MSCs because of their angiogenic and anti-inflammatory functions through paracrine secretion,^44^ and their therapeutic benefits have been proven in some clinical trials.^45, 46^ However, direct injection of MSCs suspension was reportedly ineffective in recovering ischemia due to low graft survival.^47^ Although spheroids of MSCs can improve treatments,^48^ spheroid approaches possess several drawbacks, such as low tissue adhesiveness, necrosis in spheroids, and pre-culture of MSCs. We prepared hindlimb ischemic mouse models by ligating and cutting the femoral artery (Fig. 5e). MSCs were encapsulated in the hydrogels and transplanted into ischemic tissues. CLSM observation of the transplanted MSCs revealed that µCN hydrogels retained MSCs on the target tissues and the graft survival was enhanced compared to cell suspension and non porous hydrogels (Fig. 5f). Laser Doppler imaging revealed that the ischemia occurred in the left legs post-surgery and the treatment with MSC-encapsulated µCN hydrogels gradually recovered blood perfusion, while some of non-treated mice (control) lost the limb at day 14 (Fig. 5g,h). The gross view and scoring of hindlimb ischemia showed that µCN hydrogels with MSCs improved hindlimb ischemia salvage and reduced ischemia damage compared to non-treated mice (limb loss:0% vs. 28.6%, foot loss:14.3% vs. 57.2%, toe loss:28.6% vs. 14.3%, limb rescue:57.2% vs. 0%, respectively) (Fig. 5i-k). In contrast, transplantation of the MSC suspension (MSCs alone) and non-porous hydrogels with MSCs did not improve ischemia salvage or ischemic damage. Moreover, histological observation of the hindlimb revealed that µCN hydrogels reduced immune cell infiltration, fibrosis, and muscle degeneration compared to the control, MSCs alone, and non-porous hydrogels (Fig. 5l). We expected that injectable µCN hydrogels would enhance cell migration and angiogenic signals from the material to the host tissue and supply oxygen/nutrients and cell infiltration from the host tissue to the material through porous µCN structures, which would improve material-tissue communication and therapeutic efficiency (Fig. 5m).

### Outlook

Current cell transplantation therapies are based on direct injection of cell suspensions, although delivery efficiency and graft survival are low.^6, 7^. Here, we present µCN-forming injectable porous hydrogels to improve the therapeutic efficacy of stem cell transplantation. We found that gelatin-induced LLPS can be tailored by chemical modification to form µCN structures in hydrogels and the morphologies of the porous structures. Interconnected porous structures were introduced into injectable hydrogels using an LLPS system. Porous biomaterials are considered suitable as cell-delivering scaffolds,^49^ and most of them are pre-formed types prepared by a porogen,^50–53^ foam process,^54^ and cryogelation.^55^ LLPS approaches have been also applied to fabricate pre formed porous biomaterials (e.g., polymerization of poly(2-hydroxyethylmethacrylate)^56^ and dextran-poly(ethylene glycol)^57^). However, since these methods cannot be applied for cell encapsulation due to the use of toxic materials and processes, cells need to be seeded on the scaffolds, which leads to less cell infiltration inside the scaffolds due to jamming of cells at pores.^58^ Moreover, these require additional processes, including invasive surgical treatment, fixation with suturing, and pre-culture of transplanted cells with materials in the cell processing center, which may increase the cost and time for the treatment. Injectable µCN hydrogels engineered by facile and biocompatible processes enable the homogeneous encapsulation of cells, adherence to target tissues, and delivery of cells by minimally invasive syringe injection without pre-culture, which may minimize the damage and complications associated with cell transplantation using biomaterials.

Microenvironments regulate the activity of transplanted cells, such as adhesion behavior and signal secretion, and their activities determine the response to cell transplantation therapy.^59^ Non-porous hydrogels and LLPS-induced structures such as gyroids and droplets with moderate chemical cross-linking encapsulated cells in a round morphology in the early stage of culture (24 h), whereas the µCN hydrogels promoted significant cell adhesion and spreading. The regulation of cell morphology constrained by 3D networks^60^ and cellular spreading and migration induced by biochemical and mechanostructural microenvironments in hydrogels^61^ have been previously reported; however, it is interesting that these extracellular environments can be created by the diversity of LLPS-induced structures in our system. We demonstrated that the µCN hydrogels enhanced cellular adhesion, spreading, migration, and proliferation through integrin-based mechanotransduction, where actin polymerization and the traction force associated with actomyosin-based contractility were regulated. The LLPS-induced microstructure of the hydrogel promotes ballistic and persistent cell migration and local orientation via 3D contact guidance. LLPS-induced microstructures not only modulate such cellular behaviors but are also capable of transporting oxygen and nutrients characterized by continuous cellular-level porous structures, resulting in the growth and maintenance of cells encapsulated within the hydrogels. These results indicate that µCN hydrogels allow encapsulated cells to communicate freely with the external environment of the hydrogels because of their structural features, thus potentially making them useful as injectable hydrogels for stem cell transplantation.

We utilized MSC-encapsulated injectable µCN hydrogels to induce recovery from hindlimb ischemia. In contrast to the direct injection of MSCs, µCN hydrogels can improve the graft survival of MSCs and biological communication with host tissues, such as cell infiltration and mass transport from host tissues, which may improve the long-term survival of transplants and therapeutic efficacy. Moreover, the diameter of µCN structures in hydrogels ranged around 10 µm, which would be suitable for blood vessel formation compared to macroporous structures over 150 µm.^53^ This approach that can encapsulate various cells may be extended to diverse therapies including transplantation of cardiac myocytes for severe heart failure, neurons for spinal cord injury, islets for diabetes, and T cells for adoptive cell therapy.

In addition to their application as injectable hydrogels, gelatin-based injectable hydrogels with porous structures can be used to construct various 3D-tissue structures including spheroids, fibers, and 3D-printed tissues. Since pre-gel solution of gelatin forms hydrogel when cooled below the transition temperature (approximately 30 °C), injection of pre-gel solution (37 °C) to cold PBS (4 °C) rapidly induced the gelation to temporally maintain the structures and were covalently crosslinked by thiol-ene chemistry. Spheroid tissues were constructed by droplet formation of the pregel solution in PBS (Supplementary Fig. 13a-c). µCN structures were formed in spheroids, and MSCs showed spreading in the whole hydrogel, similar to normal injection. The injection of needle formed fiber tissues and tGUPy resulted in oriented structures in the hydrogels along the flow direction (Supplementary Fig. 13d-f). The shear stress-induced orientation of µCN structures promoted the orientation of MSCs in the hydrogels, which might improve cellular functions such as proliferation.^62^ This approach has also been applied to bioprinting to engineer 3D-printed tissues (Supplementary Fig. 13g-j). Using a bioprinter, the 3D-position of the MSCs was controlled, and the printed cells showed better spreading in the hydrogels. These methods are useful for fabricating tissues with controlled 3D structures for transplantation.

Together, our findings demonstrate injectable µCN hydrogels for stem cell transplantation. This facile approach to enhance material-tissue communication with biological signals and cells through µCN structures has enormous potential as a cell-delivery carrier. These findings provide indications for the advanced design of scaffolds with controlled inner structures to contribute to tissue engineering and regenerative medicine.

## Acknowledgments

We appreciate the financial support from the Japan Society for the Promotion of Science (JSPS) KAKENHI (grant nos. 20K20207, 22H03962, and 23H01718) and the Uehara Memorial Foundation.

## Author contributions

A.N. designed and conducted this study. A.N., S.I., K.N., and H.K. performed animal experiments. A.N. and K.U. performed the mechanobiological analysis 3D-printing. A.N. wrote the manuscript. A.N., K.U., and T.T. edited the manuscript. All the authors reviewed the manuscript.

## Additional information

Supplementary Information accompanies this paper at

### Competing interests

The authors declare no competing financial interests.

**Reprints and permission information** is available online at http://npg.nature.com/reprintsandpermissions/

### Publisher’s note

Springer Nature remains neutral with regard to jurisdictional claims in published maps and institutional affiliations.

## Materials and Methods

### Materials

Porcine skin-derived gelatin (sG, *M_w_* = 180 kDa) and porcine tendon-derived gelatin (tG, *M*_w_ = 344 kDa) were purchased from Nitta Gelatin, Inc. (Osaka, Japan). 2,4,6-trinitrobenzenesulfonic acid sodium salt dihydrate (TNBS) was purchased from Tokyo Chemical Industry Co. Ltd. (Tokyo, Japan). Tris(2-carboxyethyl)phosphine hydrochloride (TCEP-HCl), collagenase, and phosphate-buffered saline (PBS) were purchased from Nacalai Tesque, Inc. (Tokyo, Japan). 5,5’-Dithiobis(2-nitrobenzoic acid) (DTNB), 2-(N-morpholino)ethanesulfonic acid (MES), RPMI1640 medium, and fetal bovine serum (FBS) were purchased from Sigma-Aldrich (St. Louis, MO, USA). Collagen casings were purchased from Nippi (Tokyo, Japan). Rhodamine-labeled bovine serum albumin, penicillin–streptomycin (P/S), trypsin, rhodamine-labeled phalloidin, anti-cTnT antibody, anti-integrin β1 antibody and 4’,6-diamidino-2-phenylindole (DAPI) were purchased from Thermo Fisher Scientific (Waltham, MA, USA). The 2-NBD-glucose was purchased from Abcam (Cambridge, UK). N-Hydroxysuccinimide (NHS) tethered Cy5.5, was purchased from Lumiprobe (Hunt Valley, MD, USA). Carprofen (RIMADYL®) was purchased from Zoetis (Florham Park, NJ, USA). The amikacin solution was purchased from Meiji Seika Pharma (Tokyo, Japan). The Human Quantikine ELISA kit for VEGF was purchased from R&D Systems (Minneapolis, MN, USA). Anti vinculin antibody was purchased from Santa Cruz Biotechnology, Inc. (USA). Dialysis membranes (molecular weight cutoff [MWCO] value:12, 000–14, 000) were purchased from Repligen (USA). Liberase and DNase I were obtained from Merck (Darmstadt, Germany). An RNA purification kit (NucleoSpin®) was purchased from TAKARA (Shiga, Japan). Dimethyl sulfoxide (DMSO), urea, iPS-derived iCell cardiomyocytes, maintenance medium, and plating medium were purchased from Fujifilm Wako (Osaka, Japan). The mouse fibroblast cell line (L929) was purchased from RIKEN (Wako, Japan). Bone marrow-derived MSC, HUVEC, NHDF, MSC growth medium, and EGM-2 cells were purchased from Lonza (Basel, Switzerland). The mouse myoblast cell line (C2C12 cell) was purchased from the European Collection of Authenticated Cell Cultures.

### Synthesis of sGTH and sGVS

For the synthesis of sGTH, 10 g of sG (amino group: 350 µmol/g) was dissolved in 160 mL of DMSO at 50 °C under stirring. The number of amino groups in SG was determined using the TNBS method. Thiobutyrolactone (714 mg, 7.0 mmol, 200 mol% equivalent to the amino groups in sG) was dispersed in 6 mL of DMSO and added to the solution. The reaction was continued for 24 h at 50 °C under stirring. To reduce the number of disulfide bonds, TCEP (1 mM) was added, and the mixture was stirred for 30 min at room temperature. The obtained solution was slowly added to 20-volumes of a cold solvent mixture of ethanol and ethyl acetate (*v*/*v*=1/1) under stirring. The precipitates were collected using a glass filter and washed three times with ethanol to remove the unreacted reagents. The precipitates collected by filtration were dried at room temperature under reduced pressure for 3 d to obtain sGTH. The degree of substitution (D.S.) of the thiol groups was calculated using the Ellman method. Briefly, sGTH was dissolved in PBS at 0.5 mg/mL of 50 °C. DTNB was added to the solution and the absorbance at 412 nm was measured using a microplate reader (Spark10M, TECAN, Switzerland). The number of thiol groups was calculated using a standard curve of *N*-acetyl cysteine.

For the synthesis of sGVS, 10 g of sGTH (thiol group: 225 µmol/g) was dissolved in 990 mL of ultra-pure water at 50 °C under stirring. Divinyl sulfone (531 mg, 4.5 mmol, 200 mol% equivalent to the thiol groups in sGTH) was dispersed in 10 mL of ultra-pure water and added to the solution. The reaction was continued for 24 h at 50 °C with stirring. TCEP (1 mM) was added and dialyzed in ultrapure water using a dialysis membrane (molecular cutoff:10 kDa) for 3 d. After freeze-drying, sGVS were obtained. Different feeding ratios (50 and 100 mol% thiobutyrolactone and divinyl sulfone) were used to synthesize sGTH and sGVS derivatives. The syntheses of sGTH and sGVS were characterized by ^1^H-nuclear magnetic resonance (^1^H-NMR, DMSO-d6, ECZ 400S, 400 MHz, JEOL, Tokyo, Japan).

### Synthesis of sG-fluorescein, sGTH-fluorescein, tGUPy-Cy5.5, and BSA-fluorescein

For the synthesis of sG-fluorescein and sGTH-fluorescein, sG and sGTH were dissolved in DMSO, and FITC (1 mol% equivalent to the amino groups in sG) was added to the solution. After precipitation, washing, and drying in the same manner as for the synthesis of sGTH, the products were further dialyzed using a dialysis membrane (molecular weight cutoff:10 kDa) for 3 d. After freeze-drying for 3 d, sG-fluorescein and sGTH-fluorescein were obtained. tGUPy (D.S.: 26, 42, 53%) was synthesized according to a previous report.^33^ in which synthesis of tGUPy-Cy5.5, tG (3 g) were dissolved in DMSO (60 mL) at 50 °C for 2 h and cooled to room temperature. NHS-tethered Cy5.5 (6.3 mg, 8.8 µmol) was added and stirred for 5 h. UPy unit (116 mg, 0.4 mmol, 45%) was then added and stirred for 24 h at room temperature. After precipitation and washing in the same manner as for the synthesis of sGTH, the products were dialyzed using a dialysis membrane (molecular weight cutoff:10 kDa) for 3 d. After freeze-drying for 3 d, tGUPy-Cy5.5 was obtained. BSA-fluorescein was dissolved in MES buffer (0.1 M, pH=6), and FITC (0.01 mol% equivalent to amino groups in BSA) was added to the solution and stirred for 24 h at room temperature. The products were then dialyzed using a dialysis membrane (molecular weight cutoff:10 kDa) for 3 d. After freeze-drying for 3 d, BSA-fluorescein was obtained.

### Preparation of µCN hydrogels

sGTH (SH groups: 225 µmol), sGVS (VS groups: 211 µmol), and tGUPy (D.S. of UPy: 42%) were dissolved in PBS (10 wt%) at 50 °C under stirring and the pH was tuned at 7.8, 6.4, and 7.4 with 1 M NaOH, respectively. The solution was maintained at 37 °C until further use. Solutions of sGTH (100 µL), sGVS (100 µL), and tGUPy (200 µL) were added and vigorously mixed using a pipette. The turbid solution was placed on a substrate and incubated for 10 min at 37 °C to obtain µCN hydrogels. To prepare non-porous hydrogels, a solution of sGTH (100 µL), sGVS (100 µL), and tG (200 µL) were used. Cross-sections of the hydrogels were observed by HE staining. The samples were fixed in a 10% formalin neutral buffer solution and stained with HE for histological observation. Images of the HE-stained sectioned gels were scanned using a digital slide scanner (NanoZoomer S210, Hamamatsu Photonics, Hamamatsu, Japan).

### CLSM observation

For visualization of LLPS, sGTH-fluorescein and tGUPy-Cy5.5 were used. The sGTH-fluorescein, sGVS, and tGUPy-Cy5.5 solutions were mixed using a pipette at a 1:1:2 volume ratio and placed on a coverslip. After the incubation for 10 min at 37 °C, the samples were observed by CLSM (CLSM 900 with Airyscan2, Zeiss, Germany). The tube length and diameter of tGUPy were analyzed using Wimasis Image Analysis (WimRetina, Spain) and ImageJ, respectively.

### Blocking test

Urea was added to tGUPy-Cy5.5 solution and dissolved at 20 M. sG-fluorescein was added to tGUPy-Cy5.5 solution at equal volume and mixed using a pipette. The final urea concentration was adjusted to 10 M. The solution was placed on a coverslip, incubated for 10 min at room temperature, and observed by CLSM.

### Rheological measurement

Rheological measurements were performed using a rheometer (MCR301; Anton Paar GmbH, Graz, Austria). sGTH, sGVS, and tG or tGUPy were mixed at 1:1:2 volume ratio at 37 °C to prepare and placed on the stage of the rheometer (pre-warmed to 37 °C) to form sGTH+sGVS+tG hydrogels (non-porous hydrogels) and sGTH+sGVS+tGUPy hydrogels (µCN hydrogels). A jig with a diameter of 10 mm was set at a gap of 1 mm. Time-dependent rheological properties were measured at 37 °C at a frequency of 10 rad/s with a 1% strain in the oscillatory mode. Rheological measurements of the hydrogels were performed before and after swelling in PBS. The hydrogels were immersed in PBS for 24 h at 37 °C and cut into a discs (10 mm) for the measurement. The angular frequency sweep measurements were performed at 37 °C at 1% strain in an oscillatory mode.

### Tensile test

Tensile strengths of the gels were measured using a tensile tester (EZ-S 500N, Shimadzu, Kyoto, Japan). A mixture of sGTH, sGVS, tG, or tGUPy was poured into a dumbbell-shaped silicone mold (total length:35 mm; width:2 mm; thickness:1 mm; ISO 37-2). After the incubation at 37 °C for 1 h, gels were released from the mold and fixed with a 1 N clump. The initial distance between clamps was 18 mm. Tensile tests were performed at a speed of 100 mm/min at 37 °C.

### Swelling ratio

The swelling ratio of the LLPS gel was measured after immersion in PBS at 37 °C. A pre-gel solution (100 µL) of non-porous and µCN hydrogels was added to a 2 mL tube. After gelation at 37 °C for 1 h, PBS (pH = 7.4) was added to each tube and incubated for 24 h at 37 °C. After incubation, the gels were collected and weighed (*W*_s_). The swollen gels were desalted by incubation in water for 24 h, freeze dried, and weighed (*W*_d_). The swelling ratio was calculated as follows:

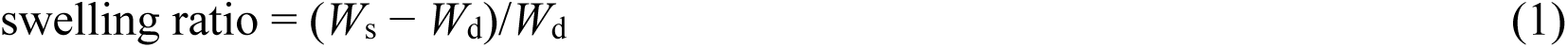

where *W*_s_ and *W*_d_ represent the weight of swelled and dried gels, respectively.

### Degradation test

A pre-gel solution (100 µL) of non-porous and µCN hydrogels was added to a 2 mL tube. After gelation at 37 °C for 1 h, PBS (1 mL) with or without 1 mg/mL of collagenase (120 U/mL) was added and incubated at 37 °C for 1, 4, 8, and 2 h. After the incubation, the supernatants were discarded, and ultrapure water was added and incubated at 25 °C for 1 h for the desalination. The resulting gel was freeze-dried and weighed.

### Tissue adhesion test

The burst strengths of the non-porous and µCN hydrogels were measured according to the American Society of Testing and Materials (ASTM) procedure (ASTM-F2392-04R, Standard Test Method for Burst Strength of Surgical Sealants). The 30 mm collagen casing discs with 3-mm pinholes were prepared. A silicone ring mold (outer and inner diameters:20 and 15 mm, respectively; thickness:1 mm) was used to fix the gel thickness. The mixture of sGTH-sGVS-tG or sGTH-sGVS-tGUPy solution (300 µL, volume ratio; 1:1:2) were added to the center of silicon mold on collagen casings and incubated for 30 min at 37 °C. The silicone mold was then removed and the samples were placed in a chamber. The burst strength (maximum pressure) was measured by running a saline solution using a syringe pump at a flow rate of 2 mL/min at 37 °C.

### Diffusion test

A mixture of sGTH-fluorescein, sGVS, and sG-Cy5.5 or sGUPy-Cy5.5 solution (20 µL, volume ratio; 1:1:2) was added to a 35 mm glass bottom dish. After incubation for 10 min, PBS (3 mL) containing rhodamine-labelled BSA (1 mg/mL) or 2-NBD-glucose (1 mg/mL) was added to each dish. The gels were observed using CLSM, and changes in the fluorescence intensity of the gels were analyzed.

### Cytocompatibility test

L929 cells were cultured in RPMI1640 medium supplemented with 10% fetal bovine serum and 1% penicillin streptomycin. sG, sGTH (−26, -39, -64), and sGVS (−25, -35, -60) were dissolved in the media at 50 °C under stirring, and the pH was tuned to 7.4 with 0.1 M NaOH. The solution was maintained at 37 °C until further use. L929 cells were seeded at 1 × 10^4^/well in a 96-well plates and cultured for 24 h at 37 °C in a 5% CO_2_ incubator. Cells were exposed to each gelatin solution and cultured for 24 h. The cell morphology was observed using an optical microscope (EVOS^®^ XL Cell Imaging System, Thermo Fisher Scientific). Cells were counted using a cell counting kit (WST-8 assay, DOJINDO, Kamishiki-gun, Japan). Briefly, 10 µL of WST-8 reagent was added to 100 µL of culture medium and incubated for 2 h. The absorbance of the medium was monitored at 450 nm using a microplate reader. The cell numbers were calculated using a standard curve. The absorbance of the cells cultured without gelatin was set at 100%.

### Encapsulation of cells in hydrogel

sGTH, sGVS, and tGUPy were dissolved in PBS (10 wt%) at 50 °C under stirring and the pH was tuned at 7.8, 6.4, and 7.4 with 1 M NaOH, respectively. sGTH, sGVS, and tGUPy were filtrated with 0.22 µm filter for sterilization. The solution was maintained at 37 °C until further use. MSCs were cultured in DMEM supplemented with 15% FBS and 1% P/S. NHDF and C2C12 cells were cultured in DMEM supplemented with 10% FBS and 1% P/S. HUVEC were cultured with EGM2-MV. iPS-derived cardiomyocytes were cultured in the maintenance medium. Cultured cells were treated with trypsin and collected as pellet after the centrifugation at 1200 rpm for 5 min. After removing the supernatant, a mixture of sGTH, sGVS, and tGUPy solution (20 µL, volume ratio: 1:1:2) was added to the pellet of each cell and mixed using a pipette. The 5 × 10^4^ cells were encapsulated in 20 µL of hydrogels for C2C12 cells, MSCs, NHDFs, and HUVECs. To prepare the cardiac tissues, 5 × 10^4^ cardiac myocytes were co cultured with 5 × 10^4^ NHDFs. For the coculture of HUVECs and MSCs, 5 × 10^4^ HUVECs and 5 × 10^4^ MSCs were used. The mixture was then placed on a chamber cover (10 × 10 mm). The samples were incubated for 10 min at 37 °C for gelation, and media (400 µL) was added and cultured for 1–2 d at 37 °C in a 5% CO_2_ incubator.

### Fluorescence staining

The cells were fixed with 4% paraformaldehyde for 30 min. After washing with PBS, the cells were permeabilized with 0.2% Triton-X for 30 min. The cells were blocked with 1% BSA/PBS for 1 h. For actin staining, cells were stained with rhodamine-labeled phalloidin (1:200) overnight at 4 °C. For cTnT staining, cells were stained with anti-cTnT antibody (1:100) for overnight at 4 °C then stained with a secondary antibody (1:400) overnight at 4 °C. After washing with PBS, cells were stained with DAPI for 1 h at 25 °C. The cellular morphology was observed using CLSM, and the area of the cells was quantified using ImageJ.

### RNA-seq analysis

MSC (1 × 10^6^ cells) were encapsulated in sGTH-sGVS-tG and sGTH-sGVS-tGUPy gels (100 µL) and cultured for 2 d. The supernatants were collected and used to quantify VEGF secretion by ELISA according to the manufacturer’s protocol. The gels were then degraded by 1.3 Wünsch U/ml Liberase and 0.2 mg/ml DNase I in serum-free DMEM for 20 min at 37 °C. Cells were collected by centrifugation at 1200 rpm for 5 min. Total RNA was extracted using an RNA microscale kit, and its concentration was determined using a Quantus Fluorometer and QuantiFluor RNA system (Promega). The RNA quality was evaluated using a 5200 Fragment Analyzer System and an Agilent HS RNA Kit (Agilent Technologies, Santa Clara, CA, USA). Libraries were constructed Using MGIEasy RNA Directional Library Prep Set (MGI Tech Co., Ltd., Kobe, Japan). The concentration of the library was analyzed using a Qubit 3.0 Fluorometer and a dsDNA HS Assay Kit (Thermo Fisher Scientific). Library quality was tested using a Fragment Analyzer and the dsDNA 915 Reagent Kit (Advanced Analytical Technologies, Santa Clara, CA USA). cDNA was prepared from the library using the MGIEasy Circularization Kit (MGI Tech Co. Ltd.), and DNA nanoballs were prepared using the DNBSEQ-G400RS High-throughput Sequencing Kit (MGI Tech Co. Ltd.. The DNA nanoballs were sequenced using DNBSEQ-G400 (2 × 100 bp). Sequence adapter, index, and primer regions were deleted using Cutadapt (ver. 1.9.91). Read sequences of ≤40 bp and low-quality sequences (of ≤Q20) were discarded using Sickle (ver. 1.33). For data analysis, read sequences were aligned using hisat2 (ver. 2.2.0). Read counts and normalization were performed using feature counts (ver. 2.0.0). The total number of reads and gene lengths among the samples were corrected using transcripts per million. GO enrichment analysis was performed using DAVID.

### *In vitro* tissue formation

MSCs were collected by trypsin treatment and centrifugation at 1200 rpm for 5 min. The 2.5 × 10^6^ cells were mixed with the pre-gel solution of sGTH, sGVS, and tGUPy (1 mL, volume ratio; 1:1:2) using a pipettor and incubated at 37 °C until the use. To form spheroid tissues, the mixed solution (20, 50, 100, 200 µL) was dropped into cold PBS (4 °C) using a pipette. To form fiber tissues, the mixed solution (1 mL) was drawn into a 1 mL syringe and injected into cold PBS (4 °C) in 100 mm dish using a PTFE needle (inner diameter: 2 mm, ∼approximately 3G). To form 3D-printed tissues, a 3D-boprinter (BIO X, Cellink, San Diego, CA, USA) was used. The mixed solution (1 mL) was filled into a syringe with a 25G needle and set in the printhead using temperature controller (37 °C). A sterilized glass dish was placed onto the stage, cooled to 4 °C. According to the program for the grid structure (3 × 3 lines, 2 × 2 cm, 0.6 cm interval, 2 layers), the solution was printed at 20 mm/s with 50 kPa (internal pump). After the fabrication process, the tissues were incubated for 30 min at room temperature for the gelation and cultured in media (20 mL) in 100 mm culture dish for 2 d at 37 °C in a 5% CO_2_ incubator.

### Biocompatibility and biodegradability test

All animal experiments were approved by the Animal Care and Use Committee of the National Institute for Materials Science (No:73-2022-03). sGTH, sGVS, and tG or tGUPy were mixed at 1:1:2 volume ratio and placed in a silicone mold with 1 mm thickness, followed by the incubation for 1 h at 37 °C. The gels formed in the mold were then dissected into 8 mm discs and subcutaneously implanted into mice. Mice (7-week-old female BALB/c nude mice (Jackson Laboratory, Bar Harbor, USA) were anesthetized by inhalation of 2% isoflurane. The backs of mice were disinfected with 70% ethanol. The gels were subcutaneously implanted into the backs of the mice. At 1, 3, 7, 14, and 28 d after implantation, the mice were euthanized by exsanguination, and tissues were collected. The samples were then fixed in 10% formalin buffer solution for 3 d and sectioned. Images of the HE-stained tissues were scanned using a digital slide scanner. The distribution of cells infiltrating the gels was quantified using the ImageJ. The edges and centers of the gels were set to 0 and 1. The distance between the cells and the edge of the gels and the cell infiltration ratio were calculated.

### Hindlimb ischemia model

Mice (7-week-old female BALB/c nude) were anesthetized by inhalation of 2% isoflurane. Hindlimb ischemia was induced as previously described.^53^ The left hindlimb was disinfected using 70% ethanol. Through a skin incision, the femoral artery and its branches were ligated using a 5-0 suture (Ethicon, Somerville, NJ, USA), and the blood vessels between the ligation points were cut. The upper and lower points of the femoral artery (the branch point of the external iliac artery and the distal point where it bifurcated into the saphenous and popliteal arteries, respectively) were ligated. After the hemostasis, MSCs (5 × 10^5^ cells/mouse) were suspended in PBS (20 µL) and encapsulated in non-porous and µCN hydrogels (20 µL) were transplanted using a pipettor. After gelation for 1 min, the skin was closed using 4-0 suture. After disinfection of the incision, the mice were intraperitoneally administered carprofen (5 mg/kg) for analgesia and the antibiotic amikacin (1 mg/kg). Blood perfusion was monitored by using a laser Doppler imaging system (OZ-2Pro, OMEGAWAVE, Tokyo, Japan). The normal (right) and ischemic limbs (left) were observed before and after treatment and on days 1, 4, 7, and 14. Mice were placed on a heating plate at 37 °C under anesthesia during the observation. The average blood flow over 6 s was recorded and compared between the ischemic and non-ischemic limbs. Ischemic damage to hindlimbs was assessed semi quantitatively. The severity of ischemic damage was determined as previously described: score 0, no change; score 1, mild discoloration; score 2, moderate/severe discoloration; score 3, necrosis; and score 4, amputation.^44, 63^ On day 14, the mice were euthanized by blood removal and the tissues were collected. The obtained tissues were fixed in 10% formalin buffer solution for 3 d, embedded in paraffin, sectioned, and stained with HE and Masson’s trichrome. The tissue images were scanned using a digital slide scanner.

### Statistical analysis

The results are expressed as mean ± SD. One-way ANOVA), followed by Tukey’s multiple comparison *post hoc* test, was used to test for differences among groups. Experiments were repeated multiple times as independent experiments. The data shown in each figure are complete datasets from representative independent experiments. None of the samples were excluded from the analysis. Statistical significance is indicated by \**P* < 0.05, \*\**P* < 0.01, \*\*\**P* < 0.001, and \*\*\*\**P* < 0.0001. Statistical analyses were performed using GraphPad Prism v.8.0 (GraphPad Software).

### Data availability

All the relevant data in this study are available from the corresponding author upon request.

