## Supporting Information for "Injectable microcapillary network hydrogels engineered by liquid-liquid phase separation for stem cell transplantation"

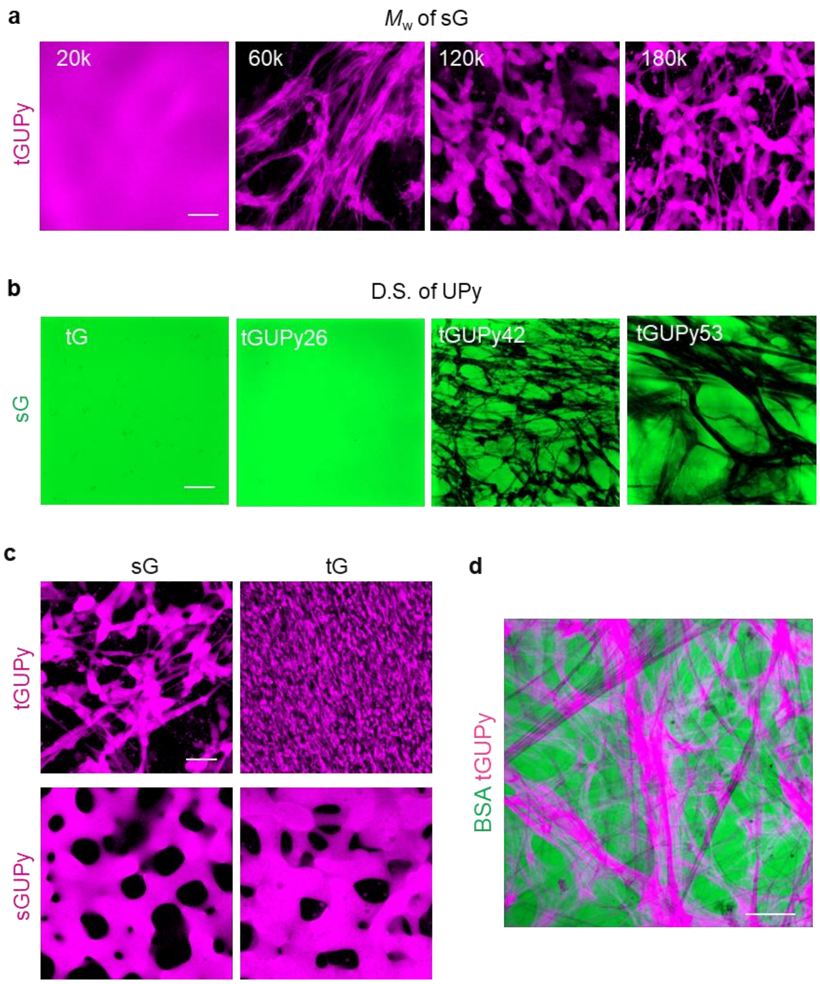


**Supplementary Fig. 1.** Effect of (**a**) molecular weight of sG and (**b**) D.S. of UPy on LLPS formation. sG-fluorescein (green) and tGUPy-Cy5.5 (violet) were used. **c**, CLSM image of LLPS structures of sG, tG, and the derivatives. tGUPy-Cy5.5 and sGUPy-Cy5.5 were used. **d**, CLSM image of LLPS structures of BSA and tGUPy. BSA-fluorescein and tGUPy-Cy5.5 were used. Scale bars represent 50 μm.

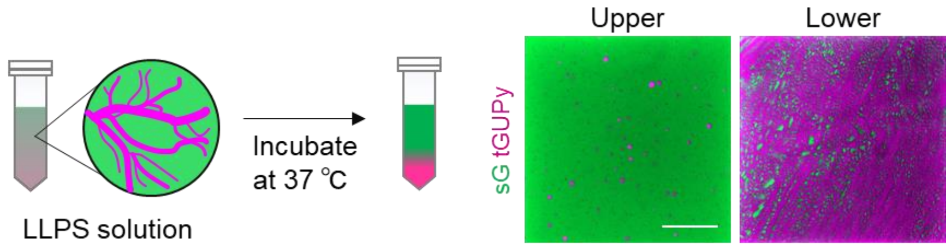


**Supplementary Fig. 2.** CLSM images of upper and lower solution of LLPS solution of sG and tGUPy after incubating at 37 °C for 1 h. sG-fluorescein and tGUPy-Cy5.5 were used. Scale bar represents 50 μm.

**Supplementary Table 1.** Synthesis of sGTH and sGVS.

| | γ-Thiobutyro-lactone (eq. to amino groups in sG[a]) | Divinylsulfone (eq. to thiol groups in sGTH) | SH[b] (μmol/g) | VS[b] (μmol/g) | D.S.[b] (%) | Molecular weight (Da) | Yield (%) | Solubility in PBS[c] |
|---|---|---|---|---|---|---|---|---|
| sG | 0 | - | 0.13 | - | 0.0 | 177000 | - | ○ |
| sGTH26 | 0.5 | - | 89 | - | 25.4 | 157000 | 87 | ○ |
| sGTH39 | 1 | - | 136 | - | 38.9 | 161000 | 79 | ○ |
| sGTH64 | 2 | - | 225 | - | 64.3 | 162000 | 79 | ○ |
| sGTH96 | 4 | - | 336 | - | 96.0 | 157000 | 82 | △ |
| sGVS25 | - | 2 | 0.59 | 88.4 | 25.3 | 152000 | 76 | ○ |
| sGVS35 | - | 2 | 14 | 122.0 | 34.9 | 170000 | 80 | ○ |
| sGVS60 | - | 2 | 14 | 211.0 | 60.3 | 172000 | 82 | ○ |

a The amino groups in as-prepared SG was calculated by determining the residual amino groups using 2,4,6,-trinitrobenzensulfonic acid (TNBS). The amounts of amino groups in sG were estimated to be 350 μmol/g. b The thiol groups in SG and degree of substitution (D.S.) of thiol and vinyl sulfone groups was calculated by Ellman method. c ○: soluble, △: partially soluble, ×: insoluble

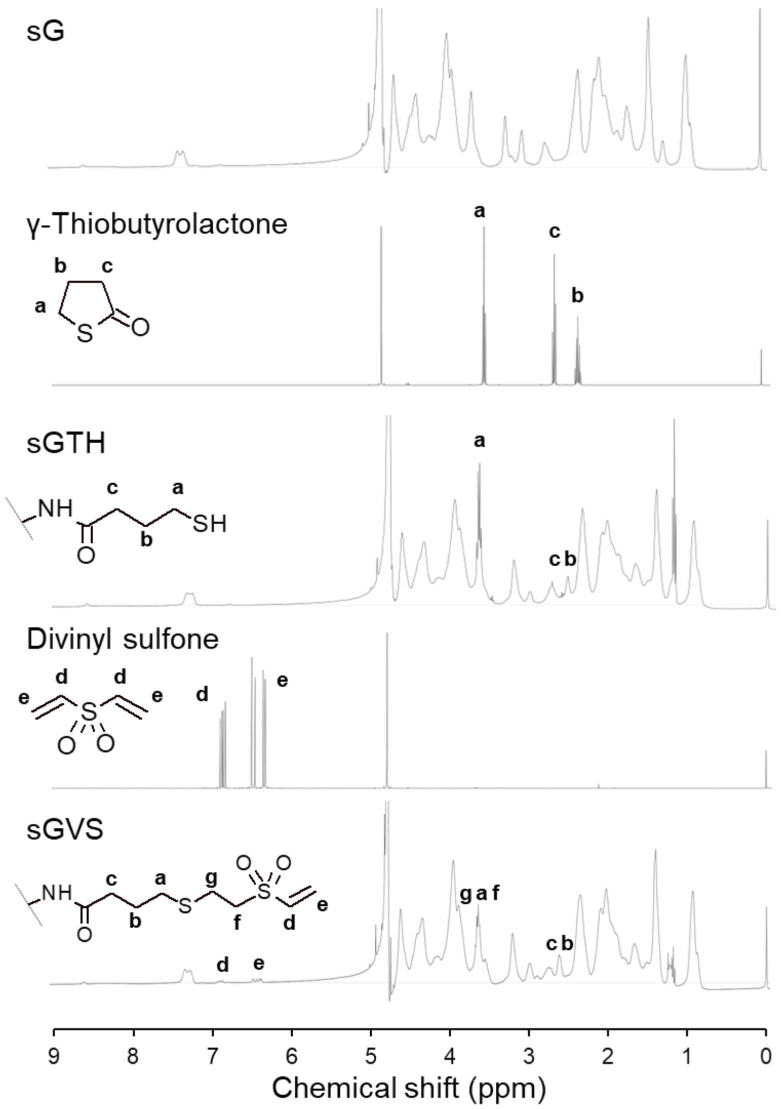


**Supplementary Fig. 3.** $^1$H-NMR spectra of sG, γ-thiobutyrolactone, sGTH, divinyl sulfone, and sGVS (400 MHz, DMSO-$d_6$).

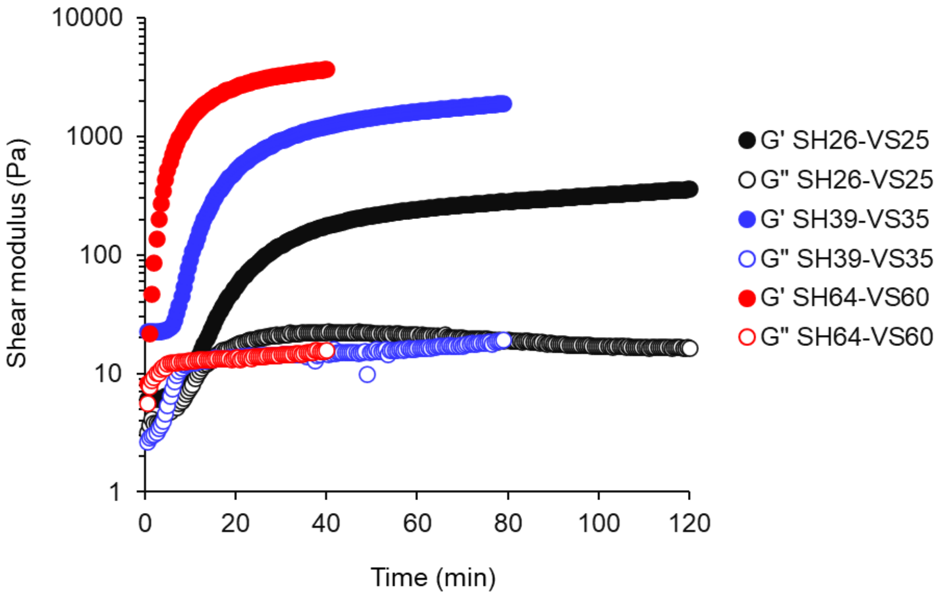


**Supplementary Fig. 4.** Rheological measurements of hydrogels of sGTH26-sGVS25 (final pH: 8.0), sGTH39-sGVS35 (final pH: 7.4), and sGTH64-sGVS60 (final pH: 7.4). The pH of sGTH26-sGVS25 was increased to 8.0 due to slow gelation. Time-dependent shear modulus changes were measured at 1% strain and 10 rad/s angular frequency at 37 $^{o}$C.

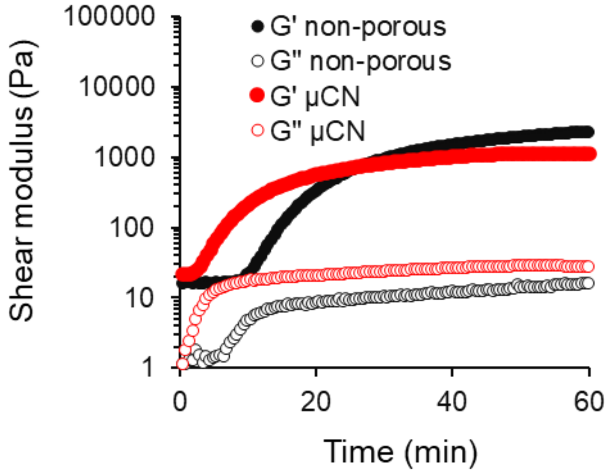


**Supplementary Fig. 5.** Rheological measurements of non-porous and μCN hydrogels. Time-dependent shear modulus changes were measured at 1% strain and 10 rad/s angular frequency at 37 $^{o}$C.

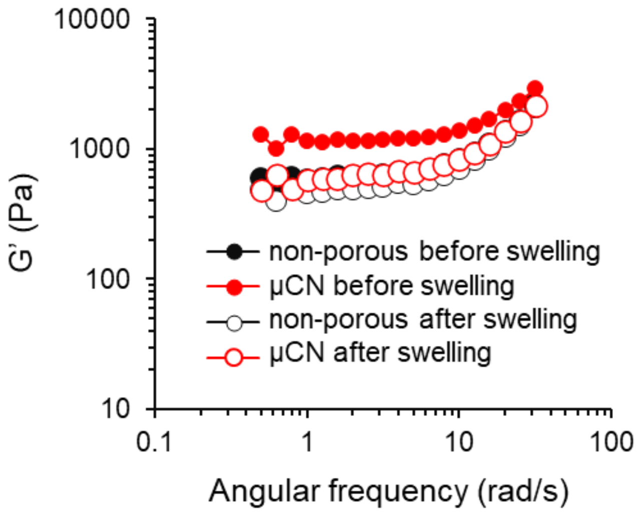


**Supplementary Fig. 6.** Rheological measurements of μCN hydrogels before and after swelling in PBS for 24 h at 37 °C. Angular frequency sweep measurements were performed at 1% strain at 37 °C.

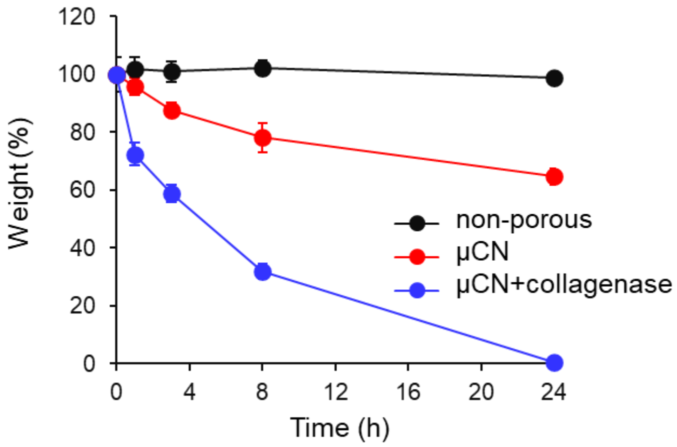


**Supplementary Fig. 7.** Weight change of non-porous (without tG) and μCN hydrogels (n = 3). Hydrogels were immersed in PBS with or without collagenase (1 mg/mL).

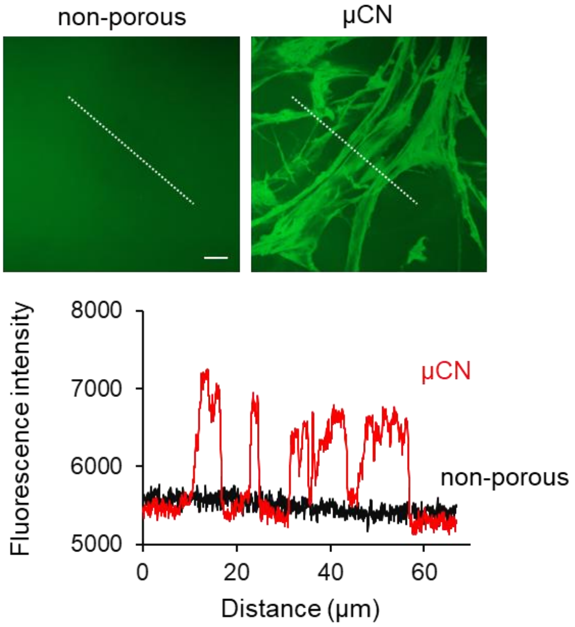


**Supplementary Fig. 8.** Diffusion test of glucose into non-porous and μCN hydrogels. Fluorescently-labeled glucose (1 mg/mL) was added to hydrogels and incubated for 1h at 37 $^{o}$C. CLSM images and line scan showed fast diffusion of glucose into gels through the lumen structures. Scale bar represents 10 μm.

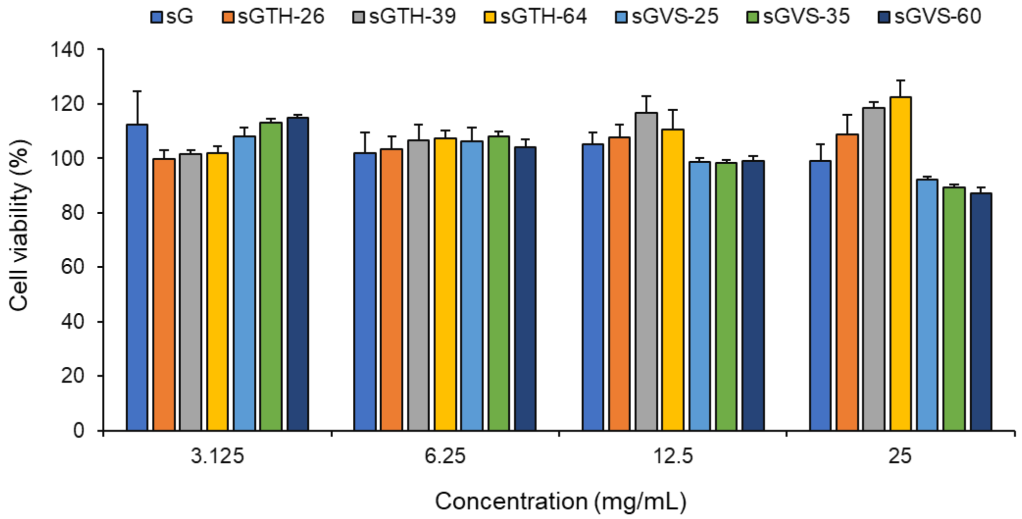


**Supplementary Fig. 9.** Cytotoxicity test of sG, sGTH, and sGVS with different D.S.. L929 cells were exposed to gelatin solution for 24 h and cell viability was checked using WST-8 assay (n=3).

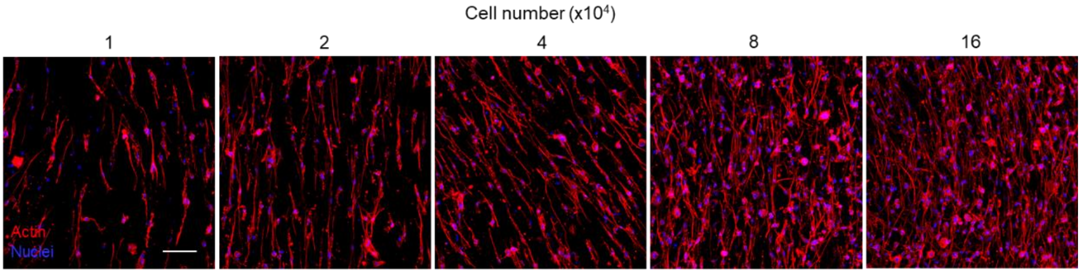


**Supplementary Fig. 10.** CLSM images of MSCs cultured in μCN hydrogels. Cells (1~16 x $10^4$ cells) were cultured in 20 μL of hydrogels for 48 h. Cells were stained with phalloidin (red) and DAPI (blue). Scale bar represents 100 μm.

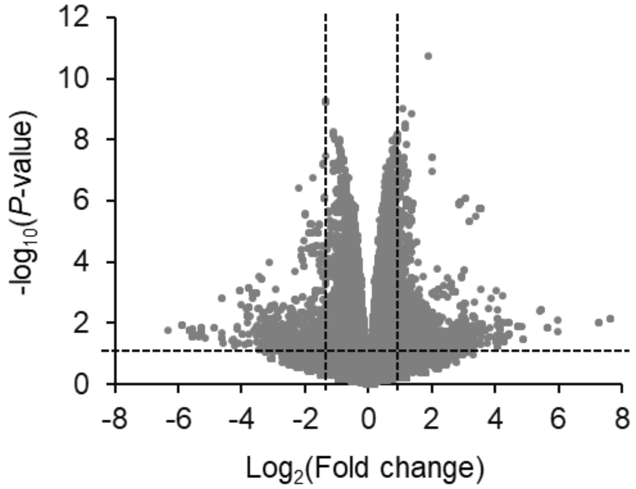


**Supplementary Fig. 11.** RNA-seq analysis of MSCs cultured in non-porous and μCN hydrogels.

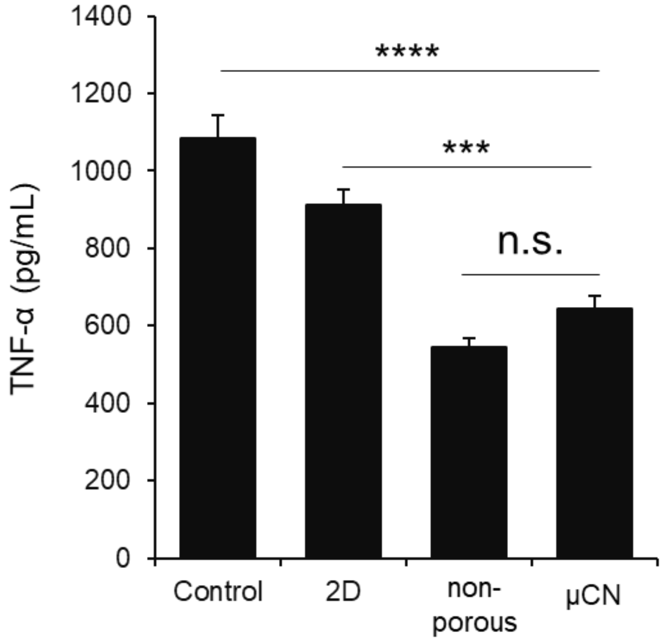


**Supplementary Fig. 12.** Secretion of TNF-α from BMDMs exposed to LPS and supernatants collected from MSCs cultured on 2D substrate and 3D-hydrogels (n=3). Data are presented as the mean ± s.d. from a representative experiment (biologically independent samples). $^{***}P < 0.001$, $^{****}P < 0.0001$ analysed by one-way ANOVA with Tukey's multiple comparison post hoc test. n.s. denotes not significant.

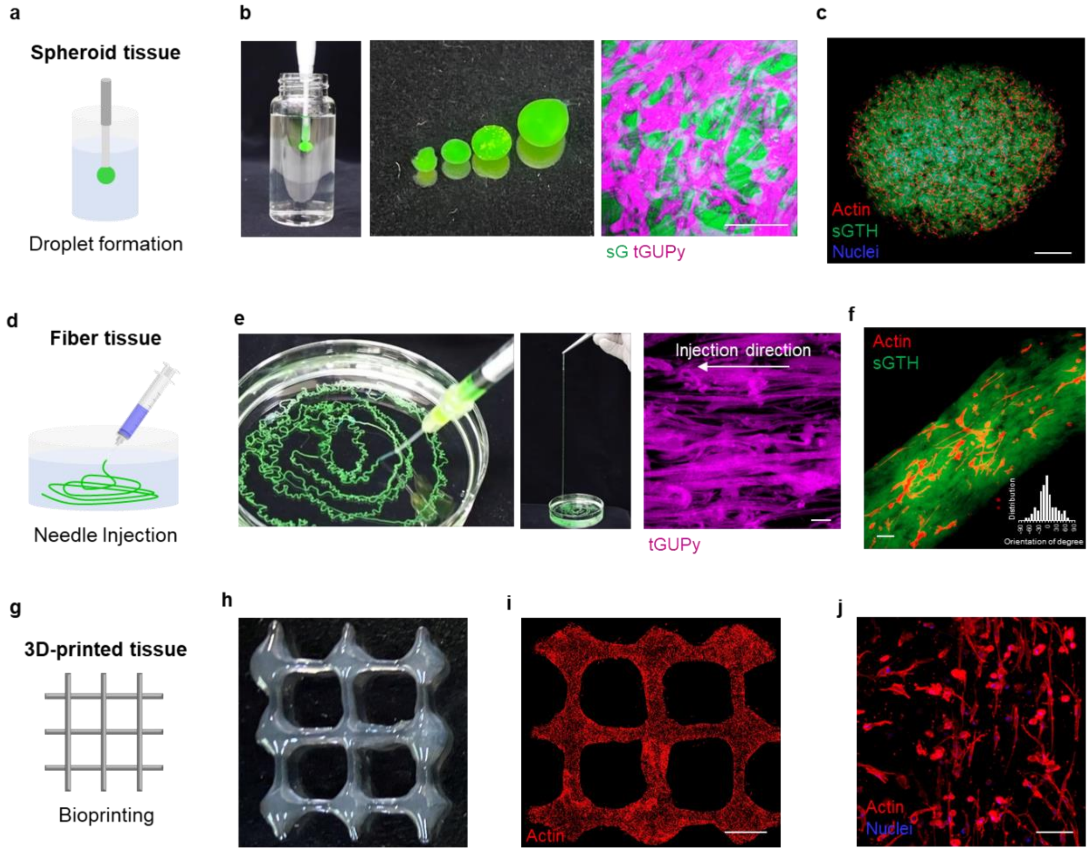


**Supplementary Fig. 13. a**,**d**,**h**, Schematic of the preparation of spheroid, fiber, and 3D-printed tissue. **b**,**e**, Photos and CLSM images of droplet-like hydrogels and fiber-like hydrogels composed of sGTH+sGVS+tGUPy. sG and tGUPy were fluorescently-labeled with fluorescein (green) and Cy5.5 (violet). **c**,**f**, CLSM images of MSC spheroids and MSC fibers. Actin and nuclei were stained with phalloidin (red) and DAPI (blue). Orientation degree in MSC fibers was quantified using ImageJ. **h**, Photo of 3D-printed hydrogels of sGTH+sGVS+tGUPy. **i**,**j**, Gross and magnified CLSM images of 3D-printed MSC tissue. Actin and nuclei were stained with phalloidin (red) and DAPI (blue). Scale bars represent 10 μm for (e), 100 μm for (b,h,j), and 500 μm for (c,i).
